# Slow Cortical Waves through Cyclicity Analysis

**DOI:** 10.1101/2021.05.16.444387

**Authors:** Ivan Abraham, Somayeh Shahsavarani, Benjamin Zimmerman, Fatima Husain, Yuliy Baryshnikov

## Abstract

Fine-grained understanding of dynamics in cortical networks is crucial in unpacking brain function. Here, we introduce a novel analytical method to characterize the dynamic interaction between distant brain regions, and apply it to data from the Human Connectome Project.

Resting-state fMRI results in time series recordings of the activity of different brain regions, which are aperiodic and lacking a base frequency. Cyclicity Analysis, a novel technique robust with respect to time-reparametrizations, is effective in recovering temporal ordering of such time series along a circular trajectory without assuming any time-scale. Our analysis detected slow cortical waves of activity propagating across the brain with consistent lead-lag relationships between specific brain regions. We also observed short bursts of task-modulated strong temporal ordering that dominate overall lead-lag relationships between pairs of regions in the brain. Our results suggest the possible role played by slow waves of information transmission between brain regions that underlie emergent cognitive function.

## 1. Introduction

The brain spontaneously generates neural activity even in the absence of any sensory inputs, motor outputs, or attention-demanding cognitive tasks. Many studies have shown that this ongoing activity is not just random noise, but carries meaningful information, reflecting a dynamic structure that can interact with perception and behavior [1, 2, 3, 4, 5, 6]. Conventionally, this structure has been characterized as synchronous patterns of neural activity in bilaterally symmetric, distant brain regions, forming brain-wide functional neural networks.

Over the past two decades, functional Magnetic Resonance Imaging (fMRI) has emerged as an important brainimaging tool to study such neural networks using a measure known as resting-state functional connectivity (FC). Studying the synchrony between spontaneous changes in fMRI blood oxygen level-dependent (BOLD) signals as a proxy for ongoing neural fluctuations provides researchers with a noninvasive tool to investigate functional organization of the brain at a network level. It is well established that resting-state FC patterns are not stationary, but rather, vary over the time course of an fMRI resting-state session [7, 8].

The temporal dynamics of spontaneous neural activity is not limited to synchronous activity. Neuroimaging studies in both humans and animal models suggest the propagation of cortical waves across the brain. For example, using electroencephalography (ECoG) recording in human subjects, Massimini and colleagues [9] showed that slow oscillations during sleep are cortical waves usually originating at anterior cortical regions. Using wide-field optical calcium imaging in mice, Matsui and colleagues [10] observed that transient neuronal coactivations were embedded within propagating waves of activity across the cortex, which was posited to carry important information underlying spatiotemporal neuronal dynamics of FC. While cortical waves of neural activity at both mesoscopic and macroscopic scales are particularly well established at high frequencies [11], much remains to be explored to better understand to what extent these cortical waves would manifest at slower frequencies measured by BOLD signals.

In the last decade, the latency structure of resting-state BOLD signals has been increasingly examined, and there is accumulating evidence for information flow even at the low frequencies sampled by the BOLD signal. In one study, Majeed and colleagues [12] found waves that moved laterally to medially, primarily in sensorimotor cortex, a result replicated in [13], using both rat and human data. More recently, Ma and Zhang [14] explored the dynamic characteristics of FC and found that transitions between resting state FC patterns follow specific sequential ordering, which is conserved in rats. Additionally, Hindricks and colleagues [15] were able to use lag-based methods to observe cortical waves in the BOLD signal in early visual cortex. Recent fMRI studies by Mitra et al. [16, 17, 18, 19] demonstrated the existence of slow restingstate inter- and intra-network propagation patterns, and the direction of such slow propagation was related to the level of arousal [17, 20]. Using lagged correlation analysis, they reported on patterns of propagated activity in resting-state BOLD signal and suggested FC synchrony as an emergent property underlying such propagation [20]. Lately, Davis and colleagues, using multielectrode array data [5], and Raut and colleagues, using fMRI and ECoG data [21], have discussed how the infra-slow spontaneous fluctuations propagating as cortical waves across the brain can be related to endogenous fluctuations in vigilance and arousal levels. Davis and colleagues suggest that cortical waves underlie changes in resting state neural networks, regulating the excitability of these networks [5]. An alternative perspective to the cortical wave was recently proposed by Huntenburg and colleagues [22] where they described how connectivity gradients predict hierarchical information flow through the cortex, and how these gradients could predict network connectivity.

Although these studies have begun to expand our understanding of the brain networks in terms of inter-regional brain interactions beyond simultaneous brain activity, developing novel tools and techniques for characterizing the mechanisms underlying such interactions still remains imperative to advance this line of research. The goal of this paper is to introduce a new tool to study such interactions, which provides fine-grained information about the temporal dynamics of BOLD signals for detecting and studying cortical waves propagating across cortical networks using resting-state fMRI data.

Previously, we applied Cyclicity Analysis [23] on restingstate fMRI data and showed that spontaneous BOLD signals comprised temporal sequences, the temporal ordering of which could be recovered along a circular trajectory [24]. In the present study, we use the data from the Human Connectome Project (HCP) [25] to reveal more complex dynamics of spontaneous BOLD signals.

## 2. Cyclicity Analysis

*Cyclicity Analysis* is a novel technique that derives pairwise temporal relations between time series using iterated path integrals (for applications in fMRI studies, see [26]). While the findings resulting from both lagged correlations and Cyclicity Analyses overlap and provide evidence in favor of the propagation of slow brain activity, they have different underlying mathematical apparatuses and assumptions, and levels of granularity. The lagged correlation method infers lag threads by deriving singular vectors of the time-delay matrix, whereas the Cyclicity Analysis method recovers inherent ordering among BOLD time series through eigenvectors of a lead matrix (a representation of the strength of temporal ordering between pairs of regions, see Section 2). Lagged correlation methods rely on interpolation and windowing to capture the dynamics of FC, which are sensitive to time delay estimation methods, autocorrelation [26], sampling variability [27], and parameters such as window length and window shift [28]. In contrast, Cyclicity Analysis offers a more robust approach with finer granularity resolution, and without assumptions regarding stationarity, state duration, or state transition.

Mathematically, the idea of Cyclicity Analysis stems from the realization that all tools based on harmonic analysis (such as Fourier transform, autocorrelation functions, power spectra, etc.) would suffer if time is reparameterized in some nonlinear fashion [23]. In physiology, one often encounters processes where essentially the same time series is playing out at different speeds at different times (think, e.g. of the heart rhythm). One then turns, naturally, to data processing tools whose results do not change if the timeline is reparameterized, or, more formally, will produce the same results on the time series related by a monotonic transformation *t* ↦ *s*(*t*), i.e. on the observations

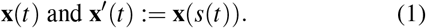

Hence, a principled data analysis tool that withstands reparametrization of the timeline should output values that remain invariant with respect to the change of parameters in (1). We will refer to such outputs, in the aggregate, as the reparamterization invariants (RI) of the trajectory **x**(*t*). The problem of finding RIs of trajectories

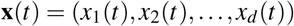

in a multidimensional space 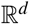 was addressed by K.-T. Chen in [29], where he established that (up to *detours*), the RIs are essentially given by *iterated integrals*. While Chen’s iterated integrals form a countable family, having a rich underlying mathematical structure, we will focus on the first nontrivial instance, the *oriented areas*: below we will show that in a broad class of models, they allow recovery of subtle interactions between the components of the time series. We note, however, that the idea of using higher order iterated integrals for data analysis was advanced by T. Lyons and his school, see e.g. [30].

### 2.1 Oriented areas

Cyclicity Analysis builds on the iterated integrals of *second order*. Such iterated integrals, not expressed in terms of the iterated integrals of first order are known as the *oriented areas*. It relies on an intuitive interpretation of the oriented area encompassed by a curve in the plane, parametrically represented by a pair of trajectories, *x* and *y*. Consider a function of time *x*(*t*) shaped as a pulse (blue curve in the Figure 1(a)) and its time shifted copy, *y*(*t*) (orange curve). We interpret the interrelationship between *x* and *y* as a *leader-follower* one: *x* leads; *y* follows. Now, to express this relationship in a reparameterization-invariant fashion, we render it using just the parametric plot (taking into account the orientation of the resulting curve in the *x* – *y* plane). The key observation now is that **when plotted against each other, these two trajectories enclose a region *R* of the plane of positive area** (as shown in Figure 1).

**Figure 1.**
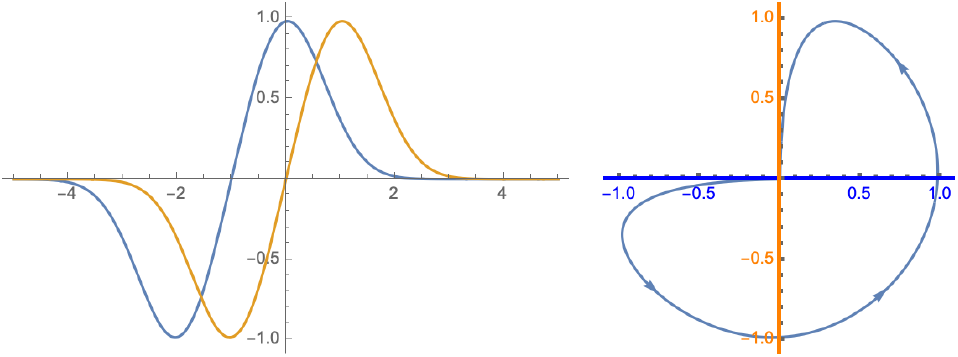
The *oriented* or *algebraic* area encompassed by a pair of time series. The left pane show a pair of time series *x* and *y* that are time-shifted copies of each other (abscissa is time). When plotted *against* each other, the right panel shows that they form a closed contour on the *x* – *y* plane.

Recall that a smooth curve partitions the plane into the open domains, such that the winding number of the curve around a point is well-defined within each of the domains (if the curve winds around a point *counterclockwise*, the point’s winding number is positive, if *clockwise*, negative). The sum of the areas of those domains, weighted by their winding numbers is called the algebraic area (encircled by the curve). Alternatively, the algebraic area can be calculated using the Green’s theorem, resulting in the representation

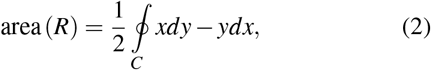

implying that the oriented area is an iterated integral of order 2 (here *C* is the curve, serving as the contour of integration, oriented by the time). Note that this area is *signed*; in particular, reversing the orientation of the curve or interchanging the variables flips the sign of the oriented area.

Note that the quantity in (2) behaves differently from the correlation coefficients. While the correlation of two signals is largest when they are proportional, the oriented areas are intrinsically antisymmetric, so that it necessarily vanishes on a pair of proportional functions. On the other hand, if the time supports of two signals do not overlap in time, their oriented area measure is also zero.

### 2.2 Lead matrices

In isolation, oriented areas just provide information about the pairwise relationships between two time series. To capture the collective information a multidimensional trajectory represents, we need to aggregate these data in a certain way.

For a trajectory in an *n*-dimensional Euclidean space 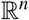, we arrange the oriented areas into the square *n* × *n* matrix (here *n* is the number of time series observed). This matrix, whose *kl*-th entry is the oriented area spanned by the pair *x_k_, x_l_* of the time series, is referred to as the *lead matrix* [23]. We remark that this matrix is obviously *skew-symmetric*, implying in particular, that its eigenvalues form pairs of purely imaginary numbers, 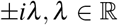, and the corresponding eigenvectors form complex-conjugated pairs with necessarily complex-valued components.

### 2.3 Chain of Offsets Model

Consider now the situation where the coordinates *x_j_, j* = 1, …, *n* of the trajectory correspond to the same function (which we will interpret here as the function on the internal clock space, a circle) 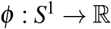, just offset by a different phase. In other words,

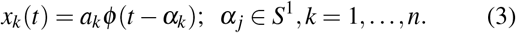

We will be referring to this model as the *Chain-Of-Offsets-Model* (COOM). The lead matrix can be readily computed in terms of the Fourier coefficients of *ϕ*: it is given by

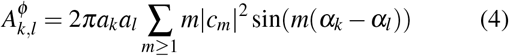

In particular, the lead matrix given by (4) decomposes into the sum of *rank two* skew symmetric matrices with entries

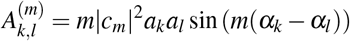

If one of the coefficients in the Fourier series for *ϕ* dominates, the skew symmetric matrix *A^ϕ^* is well approximated (in Frobenius norm) by the rank 2 matrix *A*^(*m*)^ with coefficients

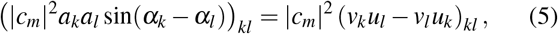

where *u_k_* = *a_k_* cos(*α_k_*), *v_k_* = *a_k_* sin(*α_k_*). In general, if a skew-symmetric operator of rank 2 is represented as *Q* = *u* ⊗ *v*′ – *v* ⊗ *u*′ (where *u, v* are linearly independent, and *v*′ denotes the conjugate vector to *v*), then one can see that *w* = −*e^−iθ^ u*/|*u*| + *v*/|*v*| is an eigenvector of *Q* with the eigenvalue *i* sin *θ*|*u*||*v*| (here *θ* is the angle between *u, v*). This implies that the real and imaginary components of the eigenvector *w_k_* = *p_k_* + *iq_k_* are obtained from the real and imaginary components *u_k_* and *v_k_* defining the matrix *A*^(*m*)^ via the linear transformation of the plane

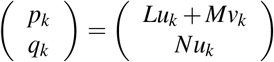

where *L* = −cos *θ*/|*u*|, *M* = 1/|*v*| and *N* = sin *θ*/|*u*|. Therefore, if

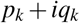

are the components of the eigenvector corresponding to the rank 2 matrix stemming from a purely harmonic COOM with the offsets *α_k_, k* = 1,…, *n*, they are just obtained from the complex numbers *a_k_* exp (*iα_k_*), *k* = 1,…, *n* by a linear transformation of the complex plane, and therefore **the cyclic order defined by the arguments of these components** is the same as **the cyclic order of the collection of points on the unit circle** {cos(*α_k_*) + *i* sin(*α_k_*)}_*k*=1,…, *n*_.

Therefore, the spectral decomposition of the lead matrix can lead to the recovery of the order in which the signals are represented by the components of the time series **x**(*t*).

## 3. Model example: Signal propagation in networks

To illustrate how Cyclicity Analysis works, we introduce a model example, where the signals are propagating through a network by gossiping. This section is dedicated to the description of a simulated model, with which we illustrate the computational pipeline of tools we apply later to the fMRI data.

Conceptually, this model, of the waves propagating in a network, broadcast from an unknown source, is a highly stylized caricature of the cortex waves. Before presenting the results on the (tentative) recovery of those waves from the fMRI readings which we do in the Section 4, it is enlightening to see what the situation is in the model example, where the ground truth is known.

### 3.1 Networks and Signals

Consider a weighted undirected graph *G* with *n* vertices, interpreted as a network, with edge lengths proportional to internode communication delays. Consider the following broadcasting protocol: if a node emits a signal, it goes to its neighbors, who will receive it after the edge-specific delay, and immediately rebroadcast it. If the signal was received by the node earlier, it ignores it. This implies that each signal reaches any given node along the shortest path connecting it to the source.

Such peer-to-peer propagation models are often referred to as *gossip, epidemic* or *first passage percolation* networks [31,32], and are relevant in studies of social networks, peer-to-peer networks, distributed resource location, etc. [33, 34, 35] Assuming such a model, one can easily recover the structure of the underlying network, *if* the shortest path lengths between the pairs of the nodes are known. Suppose a signal *s_i_* (*t*) = *f*(*t*) propagates from a *source* node *i*. If the underlying graph *G* is a connected one, then a node *j* ≠ *i* of the network will eventually receive the signal from *i* as a delayed shapeform, so that

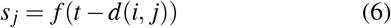

where *d*(*i, j*) is the shortest path distance between nodes *i* and *j*. For connected graphs, this distance is well-defined and finite even if there is no direct connection between nodes *i* and *j*. Quite often, one is only interested only in the ordinal information about the edgelengths (for example, the minimal spanning trees depend only on the ordering of the edgelengths). Can this ordinal information be reconstructed using *only* the observations of the (perhaps, noisy) signals at the nodes?

### 3.2 Cyclicity Analysis

We will apply the cyclicity computational pipeiline to recover the network structure from the observations of the signals *s_k_, k* = 1,…, *n*, as it manifestly fits the COOM. For each of the 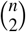 pairs of signals given by (6) observed at the nodes of the network, we form the oriented area of the corresponding 2-dimensional projection as in (2). As previously mentioned, the orientation is indicated by attaching a sign to the computed area value, with a positive sign corresponding to counterclockwise integration; interpreted as *j*(*t*) *following i*(*t*) in time.

The spectral decomposition of square matrix *A* whose entries are given by (2), i.e. the *lead matrix*, allows us to determine the relative distances of the nodes of the network to the seed node, where the signal originates. Indeed, as we argued above, the order of the arguments of the components of the (complex) eigenvector corresponding to the leading (in absolute value) eigenvalue of the lead matrix will reflect the cyclic order of the propagating wave, if the rank two skew-symmetric lead matrix corresponding to the purely harmonic signal propagating through the network is a good enough approximation of the sampled lead matrix *A*. In this case, spectral analysis of the lead matrix recovers the lag-structure between the source node and every other node.

Note that one measure of how well the assumptions and results hold is obtained by computing the eigenvalue ratio |*λ*_1_ (*A*) |/|*λ*_3_ (*A*) | of the matrix *A*, with larger ratios leading to better results. Repeating this analysis for various source nodes would then enable one to reconstruct the structure of the network.

#### 3.2.1 Model example

In Figure 2 we introduce a network on *n* = 12 nodes. The graph of Figure 2(a) is constructed by starting with the *C*_3_ graph and adding nodes with *k* = 3 edges at a time. The edges are attached to vertices at random following a distribution proportional to the vertex degree [36]. The shown matrices represent the internode distances, both as given by the edge lengths in Figure 2(b), and by the shortest path distances between pairs of nodes in Figure 2(c). (Note that the shortest distance using relays can be shorter than the direct link.)

**Figure 2.**
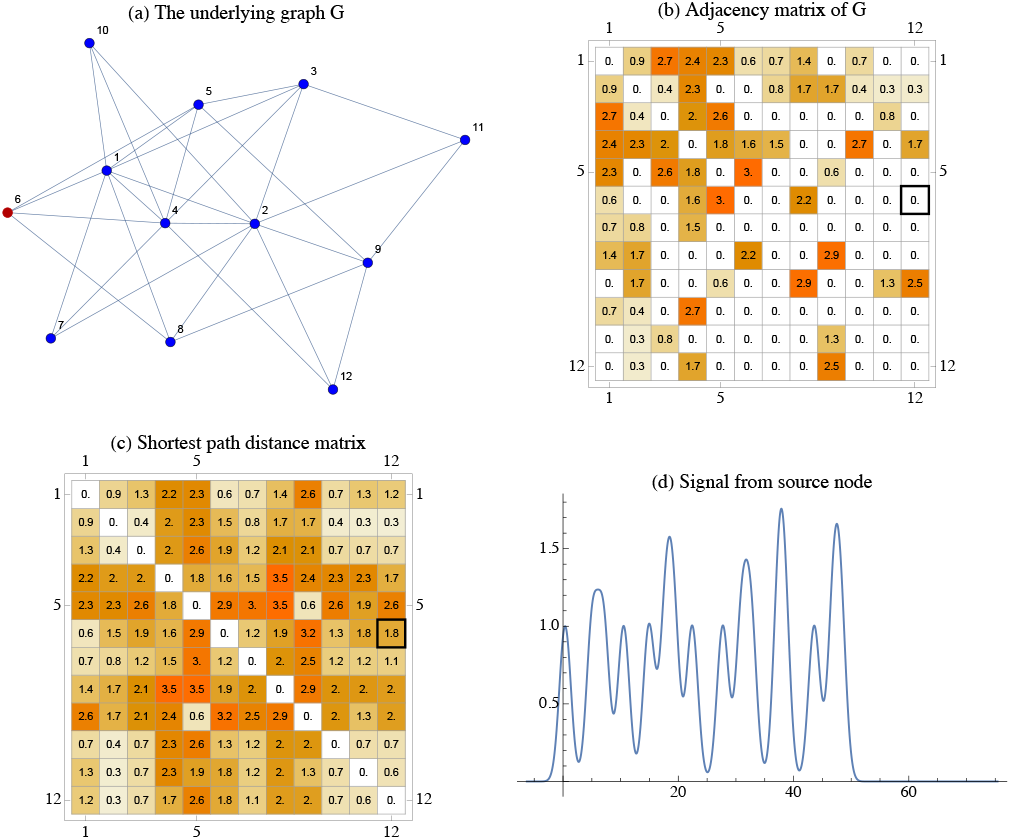
Details of the model example. **(a)** The graph *G* with *N* = 12 and source node *i* = 6, in red. **(b)** Random edge lengths. Zeros indicate the absence of a direct link between the nodes. **(c)** The shortest-path length matrix. Note that the connectivity of the graph results in all non-diagonal elements being nonzero. **(d)** The randomly generated the signal emanating from the source node.

To generate the signal in Figure 2(d) emitted by the seed note (node 6 in this case), we use a convex combination of randomly modulated Gaussian functions, randomly displaced,

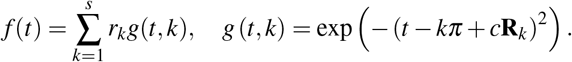

Here the displacements **R***_k_* and amplitudes *r_k_* were drawn uniformly from the appropriate intervals (implementation of the simulation is available on [a Github repository](https://gist.github.com/iabraham/3948d23417bf11e6933833b1e17ea7d4)). The signal propagates through the network according to the equation (6), to which we add a small noise, realized as the scaled Brownian motion, independently at each node.

#### 3.2.2 Recovering the network structure

The results of the cyclicity processing pipeline for the time series generated in the example of Figure 2 are presented in Figure 3. Figure 3(a) shows the sampled lead matrix. The interpretation of the entries is quite intuitive; thus one can see, for example, the vanishing oriented area between 2nd and 4th nodes; this is consistent with the fact that the network distance from the *seed* node to either of them is approximately the same. The first pair of the conjugate eigenvalues dominates in the spectrum of the lead matrix, indicative of the high resolution of the analysis as shown in Figure 3(b). The eigenvector corresponding to (one of) the leading eigenvectors has components shown in Figure 3(c). The arguments of these components, that is the angles formed by the rays pointing towards them and the ray of positive real numbers are shown on the display Figure 3(d) as the scatter plot, against the shortest distance to the seed node. The strong, essentially linear dependence between these phases and the distances to the seed shows that cyclicity can be used to detect the latter from the former, in a reparameterization invariant way.

**Figure 3.**
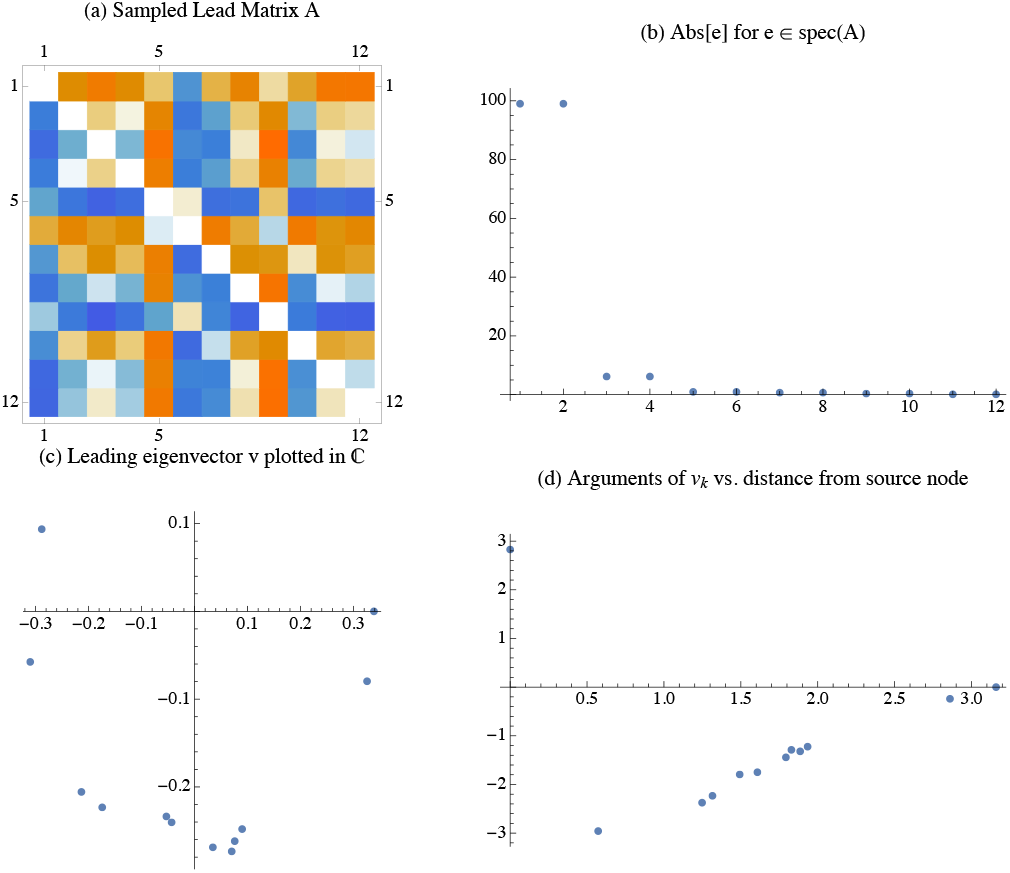
Results of the cyclicity analysis for the model example. **(a)** The skew-symmetric lead matrix. **(b)** (Absolute values of) the eigenvalues of the lead matrix. The rank 2 matrix corresponding to the first two (complex-conjugated) eigenvectors approximates the lead matrix well, as the first pair of eigenvalues dominates: |*λ*_1_| / |*λ*_3_| ≈ 20. **(c)** The “constellation” of the components of the leading eigenvector winds around the origin; the circular order of their arguments (phases) indicates the propagation of the gossip wave through the network. **(d)** More precisely, the scatter plot of these arguments against the *known* distances from the source node shows a near perfect linear dependence, implying that lag-structure is encoded with eigenvector. Typically the distances between the nodes and the source will be *unknown*, however repeated analysis with multiple sources and careful consideration of obtained complex arguments from the eigenvectors enables inference of the underlying pairwise relative distances.

#### 3.2.3 Areas dynamics

Besides recovering the ordering of the signals represented by the time series, one can extract additional information from the oriented areas, as they are accumulated in time. Namely, one can consider the integrals (2) with a variable upper bound of integration, resulting in the functions *A_ij_*(*t*), *i, j* ≤ *n*. These functions give a richer characterization of the collective behavior of the network. As a visualization tool, we will use the area gain plots, as shown in Figure 4. Since *A_ij_* = –*A_ji_* these plots are visualized in an upper triangular matrix, with a few of the plots magnified to emphasize some details.

**Figure 4.**
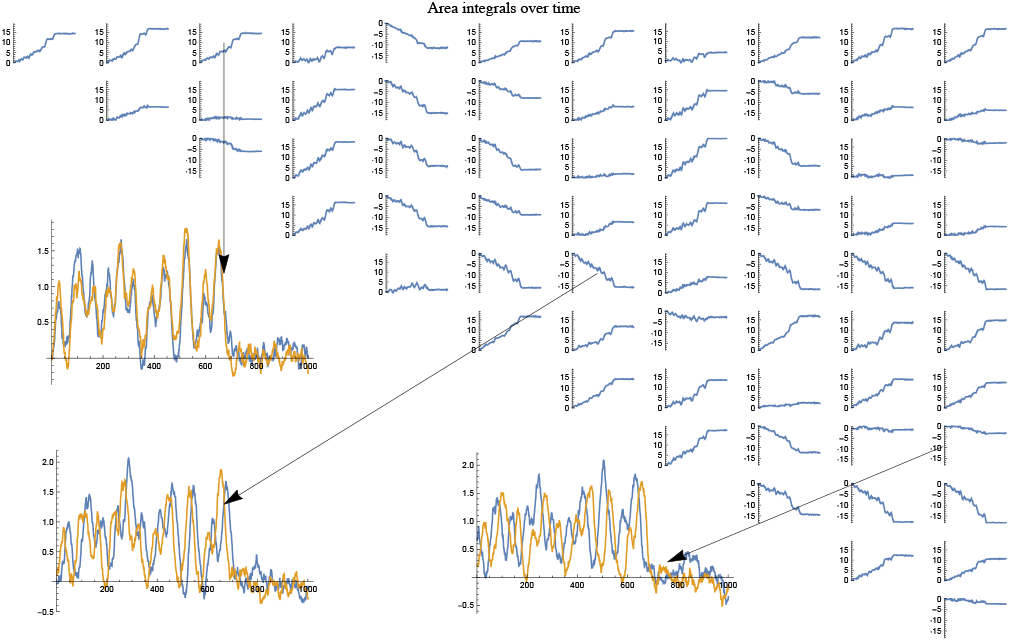
A panel showing the dynamics of the pairwise area-integrals computed as entries of the lead matrix. Since oriented areas are skew symmetric, the lower half shows insets with pairs of BOLD signals from ROIs that together generate the area dynamics. Shown are three typical instances: (top) strong directed relation as exhibited by increase, (bottom) strong directed relation as indicated by decrease and (right) a weak directed relationship as indicated my relatively flat line.

Some of the plots are flat-lines, which means there was no inferred *leader-follower* relationship between that particular pair of signals. Some plots show a sustained steady climb - *implying a strongly inferred leader-follower relationship*. The maximum gain (or drop) of the plots characterize the strength of the inferred directed relationship.

The insets in the image show the combined time series of the select pairs of signals that generated them.

## 4. Results

Now we turn to the results of Cyclicity Analysis applied to the massive dataset from the Human Connectome Project.

### 4.1 Dataset

From the Human Connectome Project 1200 Release [37, 25], we considered 889 de-noised minimally preprocessed participants who completed all the structural, resting state, and task fMRI sessions using a customized Siemens Skyra 3T scanner. Of those, 27 were excluded from the analysis for the segmentation issues noted in the [HCP Quality Control](https://wiki.humanconnectome.org/pages/viewpage.action?pageId=88901591) process or functional preprocessing errors reported in the HCP Data Release Updates. After exclusion, 862 participants remained for analysis who ranged in age from 22-45 years and included 464 females. We used the [Connectome Workbench](https://www.humanconnectome.org/software/connectome-workbench) software to extract time series from regions of interest (ROI) information in the fMRI parcellations available for download.

The ROIs extracted were 34 in number (bilateral, therefore total *N* = 68) and are listed in the Table 1. The parcellated output files were further processed in Python/R to create timecourses for the regions listed in Table 1. At a *TR* = 720 ms, for the resting state fMRI scans, this resulted in a set of arrays D, with elements of dimension 68 × 1200 that were fed into the Cyclicity Analysis. Further details of the scan protocol are available [here](https://www.humanconnectome.org/hcp-protocols). The left panel in Figure 5 shows a representative of BOLD signals in *D*.

**Table 1.** Table of 68 ROIs involved in the analysis, showing numbers for left-side regions; right-side regions run their indices as 35 through 68. Here **C** - cortex, **S** - sulcus and **P** - pole.

| ROI # | ROI full name | ROI # | ROI full name |
| --- | --- | --- | --- |
| 1 | Banks of superior temporal S. | 18 | Pars orbitalis C. |
| 2 | Caudal anterior cingulate C. | 19 | Pars triangularis C. |
| 3 | Caudal middle frontal C. | 20 | Pericalcarine C. |
| 4 | Cuneus | 21 | Postcentral C. |
| 5 | Entorhinal C. | 22 | Posterior cingulate C. |
| 6 | Fusiform C. | 23 | Precentral C. |
| 7 | Inferior parietal C. | 24 | Precuneus |
| 8 | Inferior temporal C. | 25 | Rostral anterior cingulate C. |
| 9 | Isthmus cingulate C. | 26 | Rostral middle frontal C. |
| 10 | Lateral occipital C. | 27 | Superior frontal C. |
| 11 | Lateral orbitofrontal C. | 28 | Superior parietal C. |
| 12 | Lingual C. | 29 | Superior temporal C. |
| 13 | Medial orbitofrontal C. | 30 | Supramarginal C. |
| 14 | Middle temporal C. | 31 | Frontal P. |
| 15 | Parahippocampal C. | 32 | Temporal P. |
| 16 | Paracentral C. | 33 | Transverse temporal C. |
| 17 | Pars opercularis C. | 34 | Insula |

**Figure 5.**
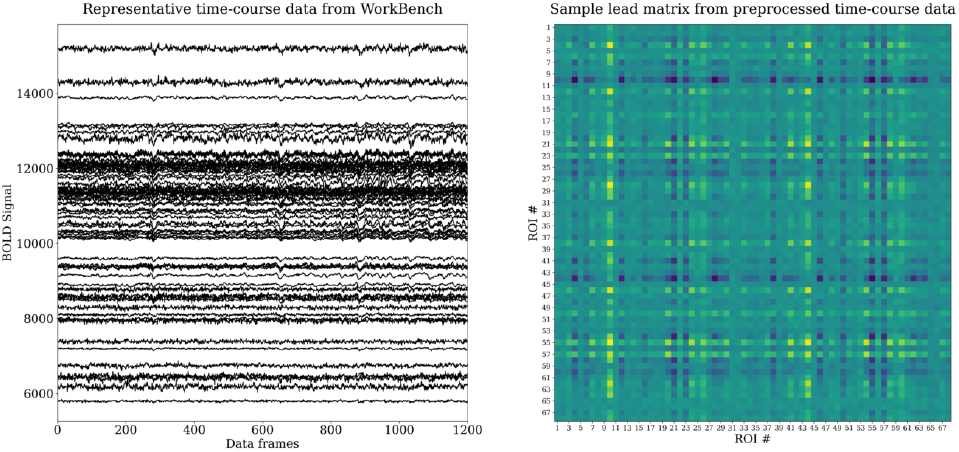
Representative time course data from the HCP dataset after processing using Connectome Workbench (left) and corresponding lead matrix (right). The lead matrix is generated after an appropriate normalization of the BOLD signal (see [24, 26]).

The HCP dataset includes fMRI scans recorded under different task paradigms. Such scans recorded during progression of certain motor and cognitve tasks were also included in part of our analysis (c.f. Section 4.6). The two task protocols are as follows - for the **motor task** [38]:

> Participants are presented with visual cues that ask them to tap their left or right fingers, squeeze their left or right toes, or move their tongue to map motor areas. Each block of a movement type lasts 12 s (10 movements), and is preceded by a 3 s cue. In each of the two runs, there are 13 blocks, with 2 of tongue movements, 4 of hand movements (2 right and 2 left), 4 of foot movements (2 right and 2 left) and three 15 s fixation blocks per run.

whereas for the **social task**[38]:

> Participants are presented with short video clips (20 s) of objects (squares, circles, triangles) either interacting in some way, or moving randomly. These videos were developed by either Castelli and colleagues (Castelli et al., 2000) or Martin and colleagues (Wheatley et al., 2007). After each video clip, participants chose between 3 possibilities: whether the objects had a social interaction (an interaction that appears as if the shapes are taking into account each others feelings and thoughts),Not Sure, or No interaction (i.e., there is no obvious interaction between the shapes and the movement appears random). Each of the two task runs has 5 video blocks (2 Mental and 3 Random in one run, 3 Mental and 2 Random in the other run) and 5 fixation blocks (15 s each).

### 4.2 Cyclicity Analysis on dataset

For Cyclicity Analysis, lead matrices were generated from the time courses and their eigenstructure analyzed. These matrices have dimension 68 × 68 with each (*i, j*) entry denoting the average leader-follower relationship between ROI *i* and ROI *j*. See the right panel in Figure 5 for a representative example of a generic lead matrix. The |*λ*_1_ |/|*λ*_3_| ratio (here *λ_k_* are eigenvalues of the lead matrix), a measure of the dominance of the rank-two approximation described in Section 2.3, was computed for all lead matrices. (Note that 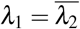 since the lead matrix is real and skew-symmetric by construction.) A subset, *S* ⊂ *D*, of the data was identified by restricting the lead matrices to have |*λ*_1_|/|*λ*_3_| ratios one standard deviation above the group average *μ_D_* (|*λ*_1_ |/|*λ*_3_|). For a visualization of the distribution of the four leading eigenvalues in *D* see the left image in Figure 6. The right image shows the distribution of |*λ*_1_ |/|*λ*_3_| ratio across *D*. We can see that *S* roughly amounts to 1/6th of all scans analyzed. The leading eigenvectors for this subset of the data was further examined. Recall that each entry *v^i^* of the eigenvector *v* of a lead matrix corresponds to one of the *N* = 68 ROIs whose time courses it was created from. Since skew-symmetric matrices only admit zero or purely imaginary eigenvalues *λ_k_* and corresponding eigenvectors *v_k_*, the elements of an eigenvector *v_k_* can be visualized as a *constellation* 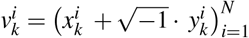, a point cloud in the complex plane with each of the *N* points representing the complex numbers 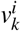, one for each ROI (essentially the scatterplot of the real versus imaginary parts of the components of the corresponding eigenvector).

**Figure 6.**
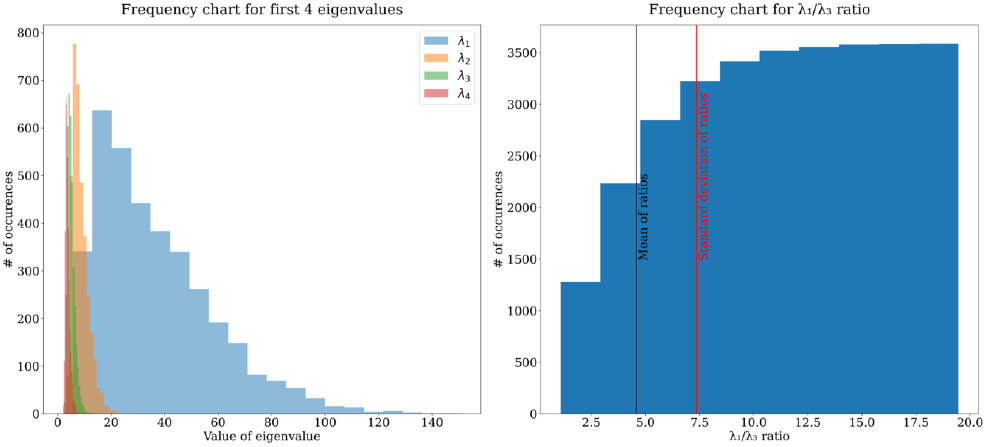
Frequency of the absolute values of four largest nonconjugate eigenvalues of the lead matrices in the data (left) along with the cumulative histogram of observed *λ*_1_/*λ*_3_ values (right). Higher *λ*_1_/*λ*_3_ values indicate more reliable outcomes from the Cyclicity Analysis pipeline.

### 4.3 Significant ROIs

In such a constellation, the points farthest from the origin correspond to ROI time courses that dominate the mode corresponding to the eigenvector *v_k_*. To identify such dominant ROIs in a principled fashion, we use the following heuristics.

Recall that in the case of a single harmonic, the constellations align along ellipses in the complex plane[23]. In the general case, the ellipse maybe perturbed, but one can argue that it still can be recovered, for example using least square regression. The best fitting ellipse defines a positive definite quadratic form on the complex plane, with its value evaluating to one on the ellipse. This form allows us to distinguish the significant points in the constellation (those where the quadratic form evaluates above a threshold, say the unity) vs. the insignificant ones (below unity, i.e. inside the ellipse). This selects a collection of ROIs dominant with respect to a particular eigenvalue. In this work we concentrate solely on the leading eigenvalues (with the largest absolute value, i.e. *λ*_1,2_ in our convention) - see Figure 7 for a representative example.

**Figure 7.**
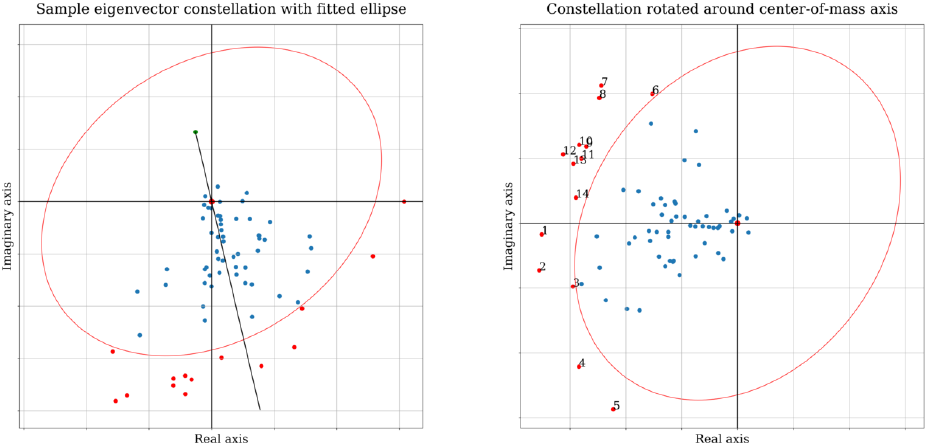
This figure shows one step in the determination of dominant ROIs in Cyclicity Analysis. The left panel shows components of the leading eigenvector visualized on the complex plane. Each point corresponds to an ROI and its BOLD signal; with greater absolute values indicating greater dominance in the multidimensional timeseries. The right panel shows the constellation of points rotated about the axis identified in the left panel to provide a consistent ordering across samples (said axis being determined by the center-of-mass for the collection of points, c.f. Section 4.5). i

This analysis resulted in the identification of a subset *R*, of 14 consistently dominant ROIs for the time courses in *S*.

### 4.4 Robustness of dominant ROIs

Since the 14 ROI subset *R* was generated by considering the restriction of the data set to *S* it important to verify that this dominance was a feature common to the entire dataset *D*. To this end, random subsets 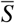 of the data of size 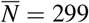 were run through the pipeline instead of *S* and the reported dominant ROIs were tracked. Figure 8 below is a visualization of this as a matrix - each row corresponds to a random subset 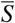 and the values along the rows represent the dominant ROIs reported from examination of 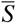 - with the color scheme set to aide visual identification of change in values between rows. The average Levenshtein distance between each pair of rows in such a matrix (across multiple trials) was found to be less than 2 illustrating that the set *R* is indeed robust with respect to the choice of the subset 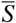.

**Figure 8.**
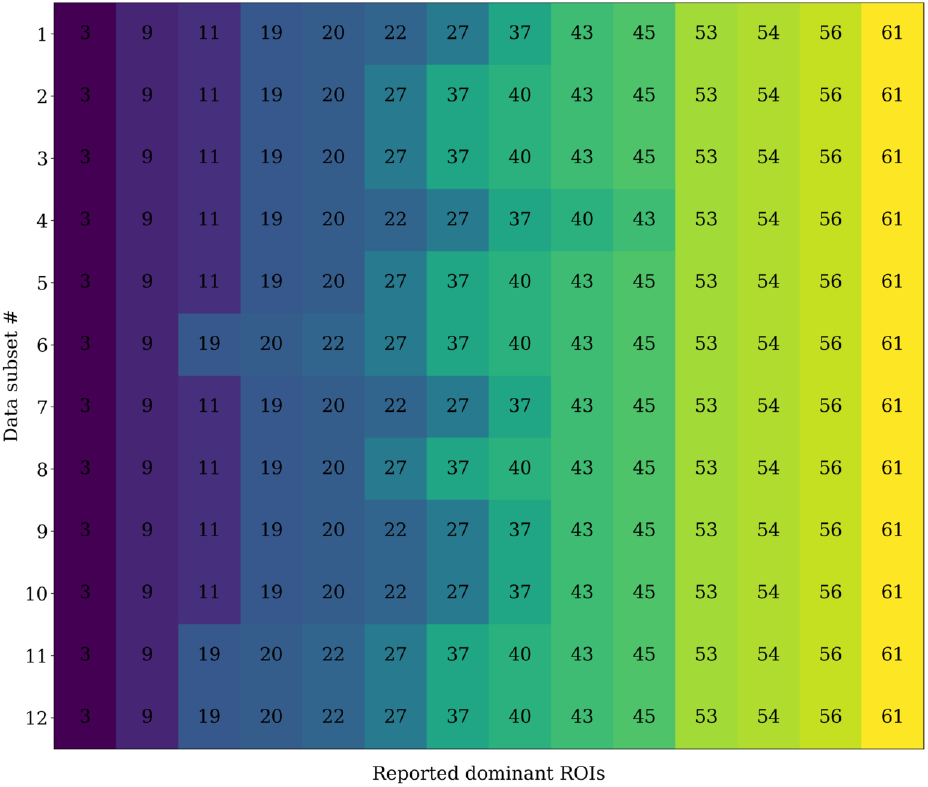
Figure showing reportedly dominant ROIs in 12 random (disjoint) partitions of the dataset. Each row of this matrix represents the most dominant ROIs reported by restricting the analysis to a random subset of the data (size *N* = 299). One can see that regardless of the subset, the same ROIs are reported to be dominant.

### 4.5 Average leader-follower relationship

As noted in [23, 24], Cyclicity Analysis is able to extract an approximate *cyclic ordering* of the phase shifts in the Chain-of-Offsets-Model from the oriented areas rendering the leader-follower relationships between pairs of time series. It is *cyclic* because the ordering of the repeating sequence of events (such as the excitation of a node) has no intrinsic starting point: only the relative positions of the phases are of import. Still, quite often there is a natural clustering, of the points in the constellation, corresponding intuitively, to a sequence of waves, coming through the network interspersed by the absence of the signal.

In this situation, one can reposition the axis corresponding to the zero phase so that it points away from the center of mass of the constellation. Choosing this ray as the abscissa, the ordering of the ROIs is then assumed to start at this axis, and proceed counterclockwise.

This is shown in Figure 7 where the left figure shows a sample constellation of the eigenvector components along with the above-mentioned axis. The right figure shows the same constellation after the standard rotation. Applying this procedure for the regions in *R* to *S* resulted in estimated cyclic orderings for these dominant ROIs. These obtained cyclic orderings can be visualized in a square *permutation count matrix* shown on Fig. 9. In this matrix the rows represent fixed ROIs and the columns represent positions in the cyclic ordering. Therefore the (*i, j*) entry of this matrix represents a value showing how many times ROI *i* showed up in position *j* in all the cyclic orderings obtained. Using this matrix, it was possible to estimate the *average cyclic ordering* for the ROIS in *R* as shown in Table 2. To do so, one first obtains a list of average positions by considering the values in each row of the matrix as samplings from a distribution and estimating their mean. Then, the permutation vector that sorts this list of mean values will correspond to the average cyclic order across the subset *S*.

**Figure 9.**
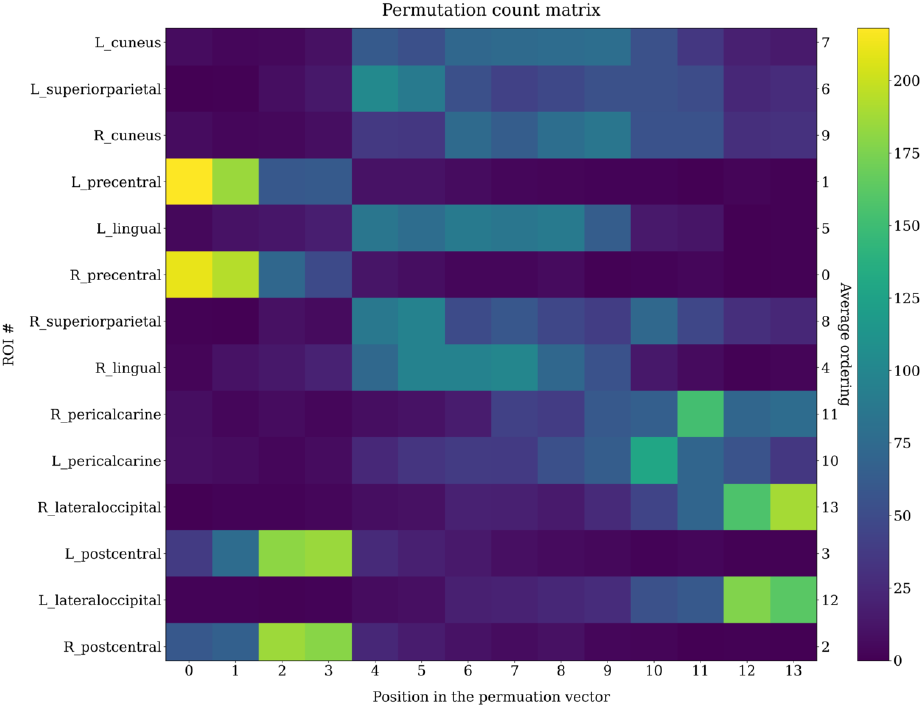
The *permutation count matrix* obtained from *S* by restricting ROIs considered to *R*. Each row of this matrix represents observed positions of the ROI in the cyclic ordering. Highlights indicate that ROI corresponding to that row *consistently* showed up in a particular position (along the columns) in obtained cyclic orderings.

**Table 2.** The fourteen dominant ROIs (including bilateral pairs) obtained and their average cyclic ordering in the dataset. The ordering shows that bilateral pairs frequently occur together in the cycle.

| ROI full name | Ordering in cycle |  |
| --- | --- | --- |
|  | Right | Left |
| *bilateral order* |  |  |
| Precentral cortex | 1 | 2 |
| Postcentral cortex | 3 | 4 |
| Lingual cortex | 5 | 6 |
| Superior parietal cortex | 9 | 7 |
| Cuneus | 10 | 8 |
| Pericalcarine | 12 | 11 |
| Lateral occipital cortex | 14 | 13 |

The stability of signal propagation across the significant ROIs in the order shown in the Table 2 is one of the main findings of this work.

### 4.6 Pairwise time-series analysis

Having established that the order in which the 14 dominating ROIs are traversed by the imputed cortical wave is a robust feature of the dataset, we further examined in depth the pairwise interactions between the time series of those 14 ROIs. For each of the 91 pairs of BOLD time series that can be formed from *R*, one can examine the area integral between the corresponding timeseries *as a function of time*, by taking the upper limit in (2) to be variable, and plotting it. Examples of such plots are shown on the Figure 10. The top-left panel shows a pair of time-series corresponding to a pair of ROIs from *R*. The top right panel shows the value of the above mentioned area integral computed between the pair of time-series signal. A consistent increasing or decreasing trend over the period of observation shows that there is an average leader (or follower) relationship between the pair of time series - i.e. activity in one region precedes (or lags) activity in the other region. While the overall trend is indicative of of this relationship, it was noted that in a large number of instances, **the increase in the value of the area integral happened in short bursts as opposed to a continuous increase**.

**Figure 10.**
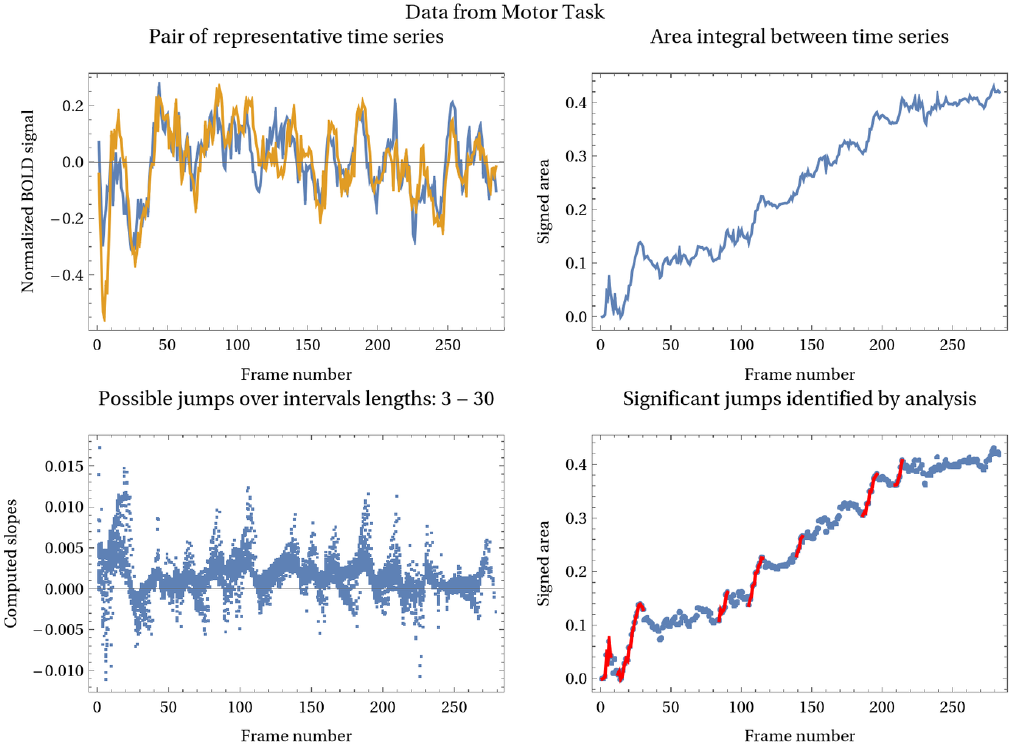
Procedure for identifying significant jumps in area integral corresponding to leader-follower activity. For each pair of timeseries as shown in the top-left panel, the area integral of Eq. (3) is computed as in the top-right figure. The slopes for possible jumps at each frame are identified and a threshold applied to designate significant jumps that contribute to the greatest overt increase in the value of the area integral.

These short periods of time where there is a significant contribution to the increase in the area integral can be termed “events” or significant periods of “directed activity” between the two pairs of brain signals. To methodically extract such instances in time, for each frame *f_k_* of the time series, successive intervals of lengths 3-30 (i.e. intervals:

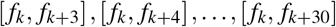

were obtained and their slope computed). These slopes obtained at each frame, are visualized in the bottom left image of Figure 10. Half of the maximum slope observed among all such intervals over all frames was taken to be a nominal threshold and *significant* jumps or contributions were deemed to be those instances when the slope of the area integral was higher than said threshold. Periods of jumps identified in such manner are shown marked in red in the bottom right image of Figure 10 and the behavior of the corresponding time series during the marked intervals shown in the inset panels of Figure 11.

**Figure 11.**
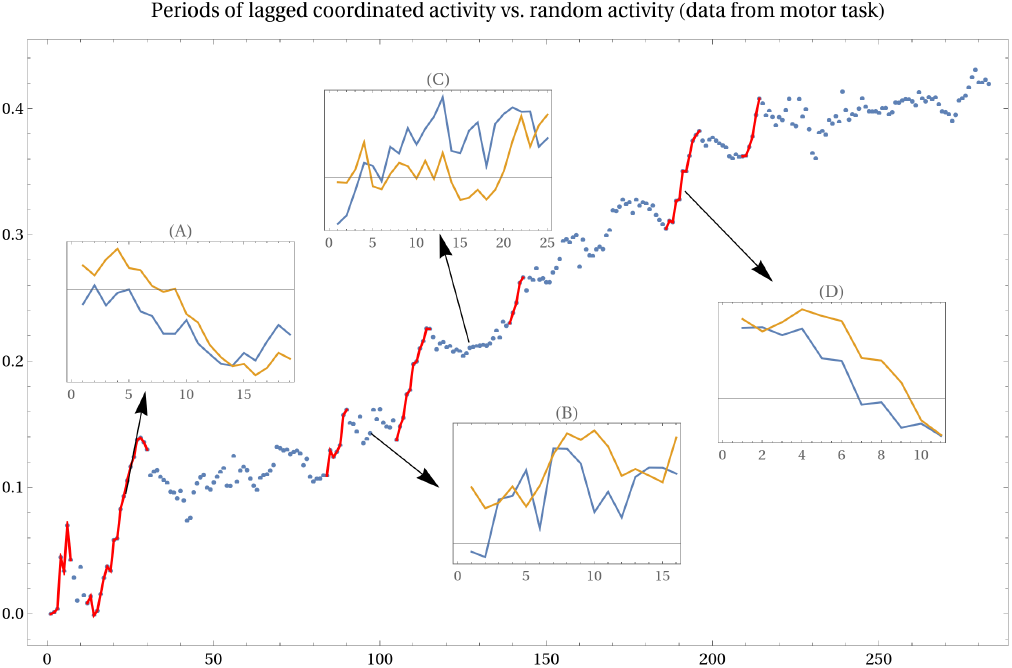
Panel showing instances of significant contribution to area integral highlighted in bottom-right image of Figure 10. Insets are titled (A) - (D) and referred to as Figure 11(A) - Figure 11(D) in the text. The inset abscissa label shows frame length of sub-interval considered from the major axis. Each pair of time-series here is the normalized BOLD signal over the titled frame. Instances when a region follows the BOLD signal in another region with a time lag correspond to greatest increases in value of the signed area integral.

The first inset Figure 11(A) corresponds to a period where there is a strong directed *leader-follower* relationship between the pair since the blue curve leads the activity of the orange one. A similar observation holds for Figure 11(D). On the other hand the relatively plateaued dynamics of the area integral shown in Figure 11(B) and 11(C) correspond to time periods between when the pair either act in concert simultaneously or a leader follower relationship is difficult to assign.

We visualized all the area integrals coming from pairs constituted from the set *R* at once by representing them in an upper-triangular array of plots. Thus, Figure 12 shows one such array. This representation allows one to capture visually, the collective trends across the pairs of time-series from *R*. For example in Figure 12, it is clear that the last two ROIs (along the columns) generally exhibit the same qualitative relationship with all other ROIs (area integral is always increasing). The insets in Figure 12 show the corresponding BOLD signals that generated the area dynamics and were specifically chosen to show *increasing, decreasing* and *plateaued* dynamics.

**Figure 12.**
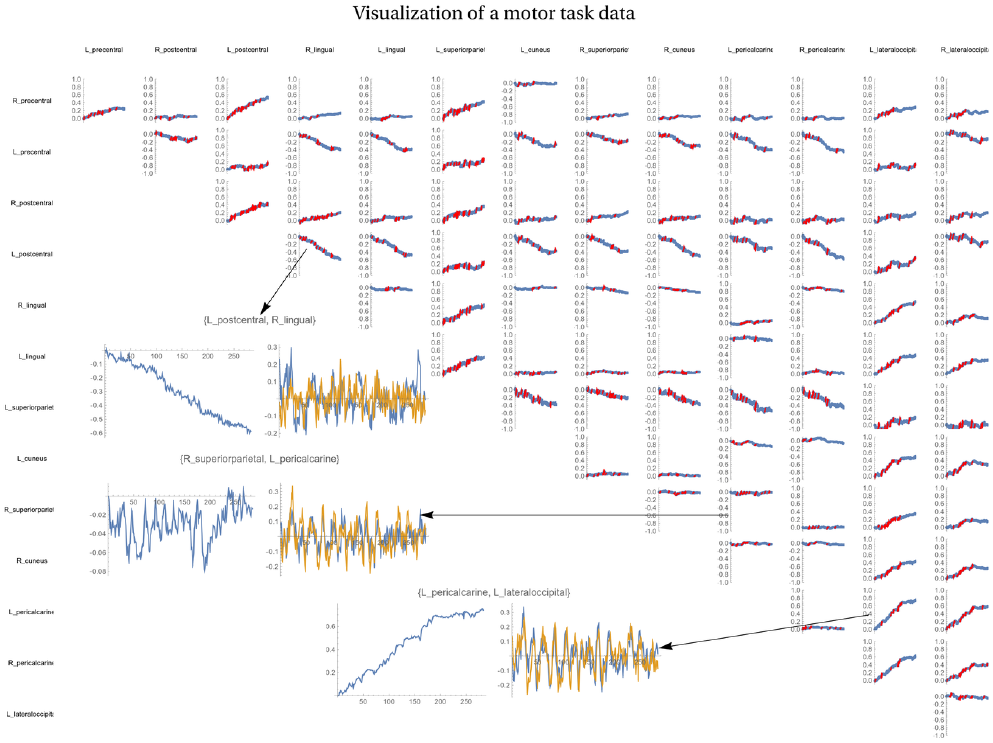
A way to visualize directed leader-follower activity between pairs of dominant ROIs obtained in the data set. A definitely increasing or decreasing trend indicates a strongly constrained directed-leader follower activity between two pairs while a variable trend indicates activity that is less constrained. In the top inset image, the blue signal leads the orange one whereas, in the bottom inset, the relationship is reversed.

While Figure 12 was generated using BOLD signals recorded during a motor-task activity, Figure 13 was generated using BOLD signals recorded during the progression of a social/cognitive task.

**Figure 13.**
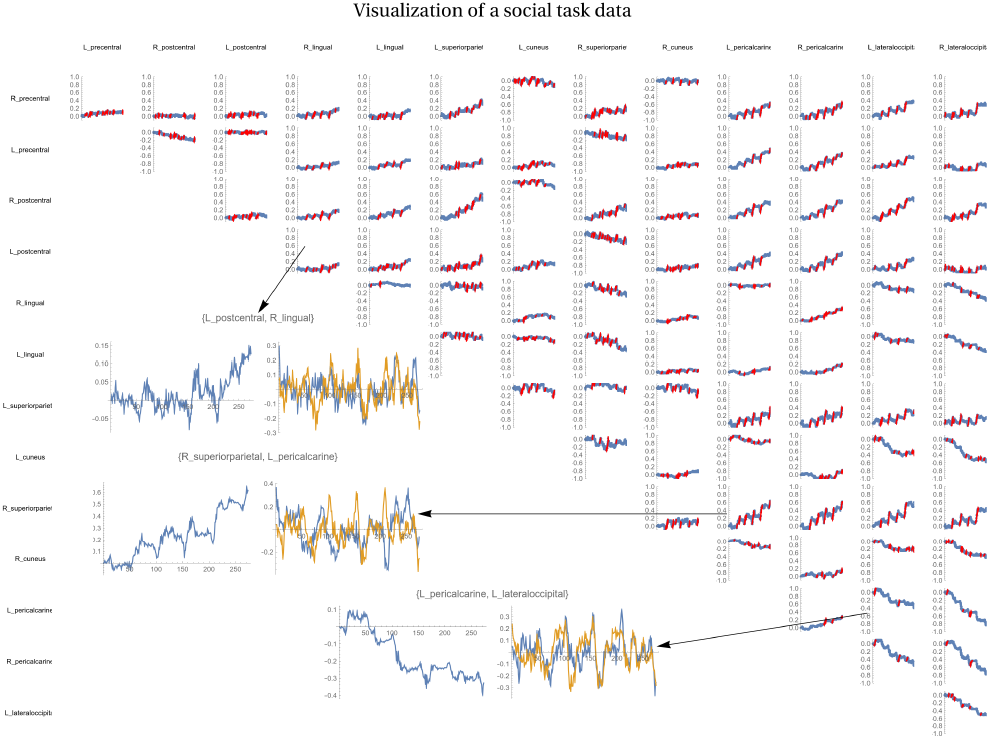
Visualization of directed activity between ROIs observed in the social cognition task. Compared to resting state or motor task scans, intermittent bursts of directed activity are a more prevalent feature in this analysis.

In Table 2 we see that precentral cortices lead the average cycle whereas lateral occitpial cortices end it. To examine the area-integral dynamics of the corresponding lead-lag pairs across different kinds of fMRI paradigms in the Connectome dataset, Figure 14 shows examples of data computed from (a) motor task, (b) a social cognition task and (c), (d) in resting state.

**Figure 14.**
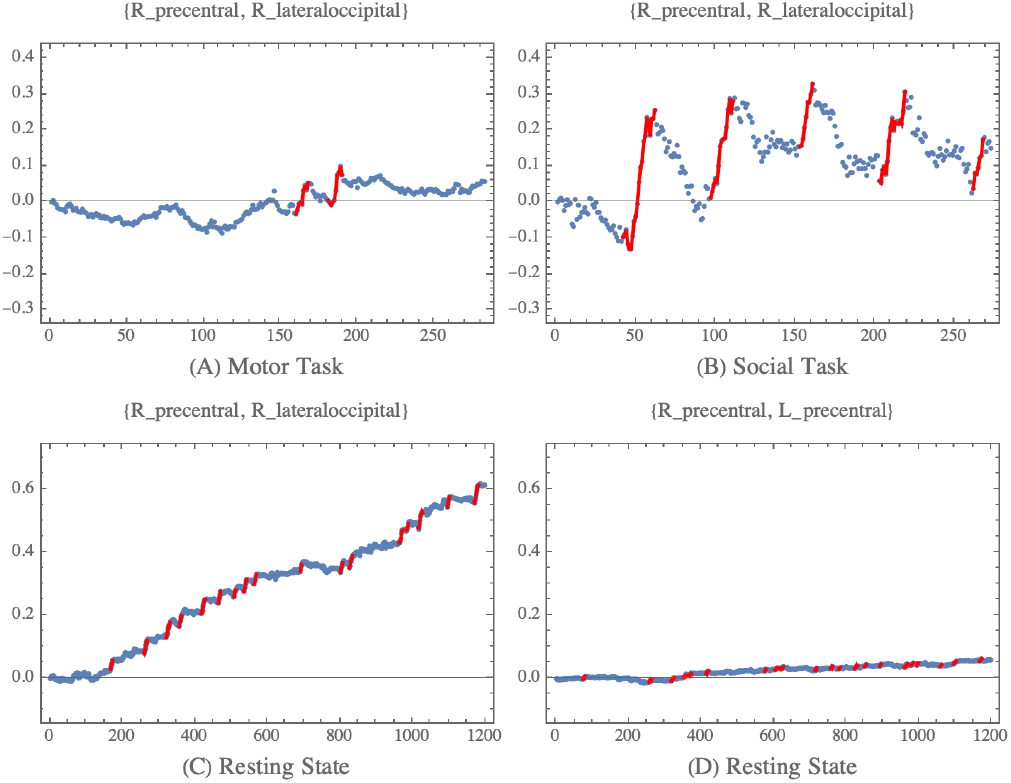
Some examples from analysis of data - **(A)** Shows the jump in the area integral for a pair of time series obtained from a motor task (`183034_r1`) and similarly, **(B)** shows jumps in a social task for the same pair in the same participant. Notice that the social task exhibits switching, i.e. there are periods of increase followed by a corresponding decrease. **(C)** shows resting state analysis (`970764_s2_r2`) while, **(D)** shows the behavior in the same motor task for the bilateral pair in the same scan.

## 5. Discussion

Cyclicity Analysis holds the potential to provide new insights into dynamics of FC, building an improved understanding of brain networks and the interplay among them. Based on this technique, we introduced an effective method in revealing the patterns of propagation for spontaneous BOLD signals across cortex. Our results provide supporting evidence for the notion of cortical waves propagating along the cortex between primary visual and transmodal areas of the brain. In addition, we found that the oriented areas between pairs of regions reflected in the lead matrix were driven mostly by short periods of sustained lead-lag relationship between the time series, suggesting a time dependent flow of information underlying the dynamic aspect of resting states hemodynamic measures.

### 5.1 Relation to Other Work

Although cortical waves have been noted at much higher frequencies, observing them in functional MRI data at low frequencies is still unexpected. At higher frequencies, cortical waves of oscillatory activity are thought to contribute to a spatiotemporal framework for neuronal communication by coordinating a range of synchronizing patterns [39, 40, 41, 42, 43, 11]. The temporal duration of these more typical cortical waves tends to be on the order of tens to hundreds of milliseconds rather than the sub-Hz frequencies to which the BOLD signal should be sensitive. Nevertheless, over the past decade, a growing body of studies has observed BOLD signal flow across cortex [13, 14, 15, 16, 21]. Taken together, these studies provide evidence that there is likely a propagation of the BOLD signal throughout the brain that dynamically carries or reflects longer periods of stable information flow between brain regions. The Cyclicity Analysis method that we have outlined here should be a useful tool for investigating these dynamics due to its invariance to the arbitrary non-linear reparameterizations of the timeline, a feature lacking in the correlation-based methods.

One of the more striking patterns our analysis uncovered was that the overall lead-lag relationships between pairs of regions in the brain were often results of short bursts of strong interactions, rather than long consistent stretches of moderate ones. This should be compared to the recent paper by Esfahlani and colleagues [44], where the authors investigated the contributions of moment-to-moment BOLD activity to the overall pattern of functional connectivity. Similar to our observation, they saw that only a small fraction of frames in the time-series explained a significant amount of the variance in the network connectivity. These fluctuations corresponded with activity patterns corresponding to fluctuations in the level of default mode and attentional control network activity, which are often viewed as in opposition to each other [45].

This opposition between the default mode and attention networks has a strong overlap with the idea of a principal gradient of macroscopic cortical organization in the brain [46]. According to this framework, a topography of connectivity patterns is reflected in the spatial distances between so-called “higher” areas of the brain, where more multi-modal, abstract, predictive information is encoded, and “lower” areas, such as the primary sensory/motor regions. This primary connectivity gradient predicts the positions of canonical resting-state networks, which are viewed in this framework as reflecting representational hierarchies rather than distinct modules. In other words, resting state networks are reflective of the temporal phase of propagating patterns, rather than as independent networks. The functions associated with various cortical networks are correlated to the level of the hierarchy of sensory or motor processing. Also, Raut et al. observed in [21] slowly propagating waves of activity measured with fMRI in humans, which were associated with cyclic fluctuations in arousal. They then replicated the result in macaques using hemisphere-wide ECoG. Critically, they found that functional networks maintained phase shifts relative to one another in their relation to autonomic arousal, rather than specific time delays. This result validates the usefulness of a time-reparametrization invariant analytical method like the one we present here.

### 5.2 Some Immediate Questions

The results of the Cyclicity Analysis, as applied to the HCP dataset, revealed some unexpected findings, that should be investigated in future studies.

The first is that the precentral and postcentral gyri, which are usually considered functionally as the primary motor and primary somatosensory cortices, sit at one end of the temporal ordering in the dominant cycle in the brain. We would expect that a cycle that reflected hierarchical processing of the visual cortex would have it’s beginning or end in some of the multi-modal processing regions of the parietal lobe that overlapped with the default mode network, such as the inferior parietal lobule or the precuneus. It is possible that there is some anatomical overlap with the postcentral gyrus, but it is not clear how the precentral gyrus would be involved. It is possible that the timing of activity in the postcentral and precentral gyri are captured in the cycle, even though the activity is not part of the same cortical wave as the hierarchical visual processing. This possibility is subtly suggested in our data, where there seem to be distinct borders between activity in visual processing areas and activity in precentral and postcentral gyri.

A second unexpected result is that the major cycle seems to end in the lateral occipital cortex rather than in the pericalcarine gyrus, where primary visual processing begins. The rest of the cycle that we observe is largely consistent with the expectations of hierarchical visual processing. The temporal ordering puts the cuneus and lingual gyri near the pericalcarine gyri and lateral occipital cortex. The superior parietal cortex activity comes near the cuneus and lingual gyri.

We showed examples of how of the time-course of lead-lag relationships changes over a run depending on what task is analyzed. In each case, we showed the lead-lag relationship between the precentral gyrus, at the starting point of the cycle we observe, and the lateral occipital cortex, which was at the end of our cyclic ordering. In the context of the resting state scan, we see strong ordering in the direction that moves in the reverse hierarchy. The precentral gyrus activity always leads and the lateral occipital cortex always follows. Rather than observing the temporal ordering between these regions as constant, it seems to move in distinct bursts, where for a few frames, the ordering is very strongly constrained. Then for some other number of frames, the ordering is relatively unconstrained. We also notice that the lead-lag relationship between these two regions takes some time to begin. It is possible that during resting state, brain activity is mostly internally generated as in mind-wandering. This state of mind-wandering takes some time to start, but once initiated, since there is no stimulus to attend to, activity mostly occurs in the higher-to-lower direction in the hierarchy reflecting endogenous processing.

In the context of the social task (see Figure 13), we observed a strikingly different pattern. This task consisted of a 5 video blocks where there was a period of watching a video and then judging whether or not the objects in the video appear to perceive the other object’s feelings and thoughts. In this task, we still observe strong directional constraint, but it seems to shift in which area is the leader and which is the follower. This may be interpreted as periods of time that switch between internally generated (endogenous) activity and externally generated (exogenous) activity, that may correspond with periods of watching the videos and periods of making judgements of the objects’ intentions within the video.

We observe a less interpretable pattern of activity in the context of a motor task. Here the participants are presented with 3-second cues to do 12-second motor movements. There are 13 motor blocks and three 15 second fixation blocks in each run. Like the motor task, we observe switches between the which region is the leader and follower during the run. However, it is difficult to match the activity observed to events in the task. This may be because the cuing events and motor movement events are too short to be adequately captured in the low frequency movement across the brain. We may be observing some aliasing in the directionality of the lead-lag signal. It is also possible that in the motor task, there is more separability between activity belonging to hierarchical visual processing and motor processing. Both of these hypotheses are somewhat supported by the observation that if we observe purely visual regions, which are close together in both anatomy and expected temporal hierarchy, such as the pericalcarine gyrus and lateral occipital gyrus, we observe a very different pattern of temporal ordering. There are still alternations in the direction of temporal order, but they are more frequent and weaker in magnitude. Perhaps areas that are closer together are better able to capture the short time periods between subsequent cues and motor events, or perhaps it reflects less contamination from motor activity coming from the precentral gyrus.

### 5.3 Remarks & Caveats

Our analysis identified a temporal ordering that propagated from the primary occipital regions towards more transmodal regions. However, it should be noted that our analysis focused on analyzing the first eigenvector of the lead matrix, and we are specifically showing patterns of activity from subjects selected based on having high |*λ*_1_/*λ*_3_| ratios in the lead matrices. This choice likely selects for subjects with strong leader-follower activity along a single direction. It is possible that analysis of the other eigenvectors would reveal a pattern of temporal ordering moving in the other directions, reflecting alternative gradients of direction from regions higher in the cortical hierarchy to the primary sensory regions lower in the cortical hierarchy or more local interactions, such as the signaling across hemispheres.

## 6. Conclusion

In this paper, we deployed Cyclicity Analysis to detect transient states in the brain. This new tool complements recent advancements in effective or directional connectivity research.

The method outlined here has intrinsic advantages due to its lack of assumptions regarding the temporal properties of the BOLD signal dynamics. It does not assume stationarity of the time-series or specific properties of latency, state duration, or state transitions, which could bias correlational, spectral, or lag-based approaches. We have shown in group data a primary propagating wave of BOLD activity during resting state from somatomotor cortex to early visual cortex. We further observed that these propagating waves appear to switch direction in a task-dependent manner from examining individual subjects.

The results of the data analysis presented here open a wide range of future research directions. Future work should apply Cyclicity Analysis techniques to other measures of human brain activity that are more reflective of direct neural activity, including electroencephalography (EEG), electrocorticography (ECoG), the fast optical signal, and magnetoencephalography (MEG). Applying Cyclicity Analysis to multiple sensing modalities may help clarify the relationship between patterns observed in the fast temporal domain of neuronal activation and the longer duration patterns observed in the BOLD signal, which may be more reflective of broader stable states of whole-brain function.

## Acknowledgments

Data were provided, in part, by the Human Connectome Project, WU-Minn Consortium (Principal Investigators: David Van Essen and Kamil Ugurbil; 1U54MH091657) funded by the 16 NIH Institutes and Centers that support the NIH Blueprint for Neuroscience Research; and by the McDonnell Center for Systems Neuroscience at Washington University.

The authors are grateful to Prof. Marcus Raichle for making available an early text of [21].

YB was partially supported by the ARO, through MURI SLICE.

